# Environmental RNA/DNA Metabarcoding for Ecological Risk Assessment of Chemicals Based on Response of Benthic Communities in Natural Environments

**DOI:** 10.1101/2025.05.29.656937

**Authors:** Yasuaki Inoue, Kaede Miyata, Masayuki Yamane, Hiroshi Honda

## Abstract

To achieve the current Nature Positive goals, it is important to identify the effects of chemicals on ecosystems and strategically reduce their risks. We therefore estimated the safe concentration (SC) of linear alkylbenzene sulfonate (LAS) by investigating the relationship between benthic communities detected via environmental RNA (eRNA) and environmental DNA (eDNA) metabarcoding and the water quality in a river contaminated with LAS. Non-metric multidimensional scaling of benthic communities detected via eRNA indicated significant differences between contaminated and non-contaminated sites, and redundancy analysis showed significant correlation between the benthic communities and LAS concentration. These results demonstrate that eRNA is more sensitive than eDNA for capturing the community changes that result from LAS contamination. No significant differences were observed in biodiversity indices between the highest LAS concentration site and non-contaminated site; thus, the SC was estimated at >47 µg/L. The results suggest that, although hazard assessment using the assessment factor (AF) leads to conservative estimation of the effects of chemicals because the predicted no-effect concentration in previous studies is lower than the SC, hazard assessment based on the response of biological communities using eRNA metabarcoding can refine such ecological assessments, developing the environmental management standards to achieve nature-positive goals.

## 1. INTRODUCTION

The decline in the ecological health of aquatic environments is a major challenge for human society^1,2^. At the 15th Conference of Parties for the Convention on Biological Diversity (COP15) in 2022, an urgent call for action to halt and reverse biodiversity loss and allow nature to recover was set as a goal for 2030^3,4^. The latest planetary boundary assessment reported that anthropogenic chemicals are already above safe levels^5^, it is important to monitor their impact on the ecosystem and establish methodologies that can strategically reduce the risks and realize the Nature Positive goals^6^.

Ecological risk assessment, in the form of exposure and hazard assessment, can assess the degree to which chemicals negatively affect ecosystems^7–10^. The most common method predicts the effect of chemicals on ecosystems by dividing toxicity data by an assessment factor (AF) to calculate the predicted no-effect concentration (PNEC) below which no adverse effects are expected^10,11^. Contaminated aquatic ecosystems are affected by both direct toxicity and indirect changes in ecological interaction; thus, community-level assessments can provide clearer indication of the effects of chemicals^12–15^. Community-level assessments while considering complex ecological interactions contribute not only to assessing the effects of chemicals on real ecosystem, but also validating the hazard assessments by AF method^16^.

Environmental nucleic acid (eNA; i.e., environmental RNA (eRNA) and environmental DNA (eDNA)) metabarcoding is an alternative biodiversity monitoring technique^17–20^, and can estimate the biological composition of an environment via noninvasive biological survey^21–23^. ERNA reduces false positives because it is less stable than eDNA, allowing for the accurate prediction of biological communities^24–27^. We hypothesized that the high sensitivity of eNA in capturing the biological communities in aquatic environments contribute to evaluate community-level effects and estimate the safe concentration (SC) of chemicals within an ecosystem. To the best of our knowledge, no hazard assessment studies have as yet used eNA metabarcoding to investigate the response of community structures to chemical contamination^28,29^.

Herein, the effects of linear alkylbenzene sulfonate (LAS) contamination on an ecosystem were investigated by analyzing the relationships between biological communities, detected by eNA metabarcoding and traditional field surveys (TFS), and water quality parameters at sampling sites in a LAS-contaminated mountain stream. The SC was estimated by comparing various biodiversity indices at different sampling sites with varying LAS concentrations. LAS is a major anionic surfactant that is used in detergents worldwide and is closely related to human activity. Although numerous laboratory-based toxicity tests have been conducted, the effects of LAS on environmental ecosystems have rarely been studied. Benthic macroinvertebrates are suitable indicators of water quality because they are relatively sedentary and sensitive to changes in water quality and environmental health^30^. The main goals of this study were to (1) clarify the utility of eNA for community-level hazard assessment, (2) evaluate whether eRNA can more sensitively capture the effects of chemical contamination than eDNA, and (3) estimate the SC for LAS to protect benthic biodiversity in rivers and identify issues in hazard assessment using the AF method.

## 2. MATERIALS AND METHODS

### 2.1. Sampling Site

A river in the Kanto region of Japan was selected as the study site (Figure 1a). The Japanese pollutant release and transfer register (PRTR) system^31,32^ indicates that a nearby plant discharges only LAS into the river; thus, it was expected that the area downstream would be contaminated with LAS. To minimize the influence of factors other than LAS, sampling was also performed at points with similar environmental factors, such as river width and bottom sediment size. River water samples were collected from three sites: one 350 m upstream of the chemical plant effluent outfall ("upstream site”), one 50 m downstream of the outfall ("outfall site”), and one 1700 m downstream of the outfall ("downstream site”) (Figure 1b). Field surveys using TFS and eRNA/DNA metabarcoding and water quality measurements were conducted at all three sites. Sampling was conducted with permission from the city office and the local fishery cooperative association. Water quality measurements and eNA metabarcoding were conducted six times at each site and TFS once (upstream site: December 14, outfall site: December 11, and downstream site: December 13, 2023) from December 2023 to February 2024 (Table S1).

**Figure 1.**
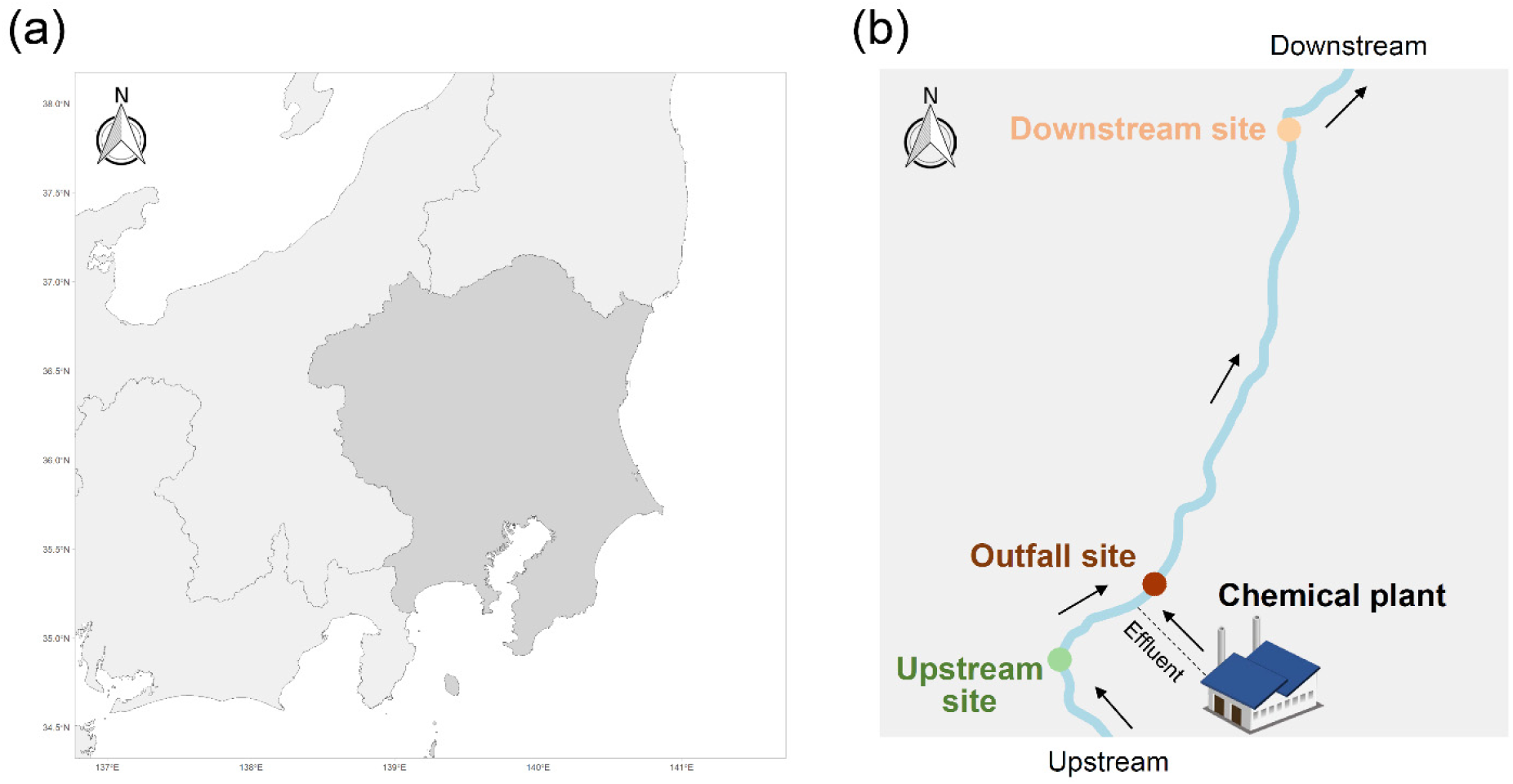
Geographical location. (a) Kanto region of Japan. (b) Locations at which chemical plant effluents are discharged into the river and locations from which samples were collected. The upstream site was approximately 350 m upstream of the effluent outfall, while the outfall and downstream sites were approximately 50 and 1700 m downstream of the effluent outfall, respectively.

### 2.2. Water quality and LAS concentration analyses

Seven water quality parameters were measured to confirm the similarity in the physicochemical characteristics at the different sampling sites. Dissolved oxygen (DO) and pH were measured in situ using a pH/DO meter D-210PD-S (HORIBA, Kyoto, Japan); the five-day biochemical oxygen demand (BOD) was measured using BOD dilution method (JIS 0102 21 and 32.2); suspended solids (SS) were measured via filtration method with glass fiber filter paper (JIS K 0102 14.1); *Escherichia coli* (*E. coli*) numbers were determined using the method specified by the Ministry of the Environment, Government of Japan^33^; LAS concentration was determined using the method described in the Supplementary Information (Text S1); and total organic carbon (TOC) was determined using the combustion oxidation method (JIS 0102 22.1). Measured values that were below the lower limit of detection (LOD) were replaced with half the LOD value.

### 2.3. Water sampling for eRNA/DNA metabarcoding analysis

River water samples (20 L) were collected from each site and stored on ice. Filtration was performed in the field using Sterivex cartridges (0.45 μm; Millipore SVHV010RS; Merck, Tokyo, Japan) with a flow-through pump. Five duplicate samples were collected from all sites on each sampling day and prepared by filtering 3 L of river water for eRNA/eDNA analysis, with a filtration blank prepared at each sampling site by filtering 3 L of distilled water using the same method. To prevent nucleic acid degradation, 2 mL of RNAlater was added to the Sterivex cartridges and the resulting samples stored at -20 °C until nucleic acid extraction could be performed.

### 2.4. ERNA/eDNA extraction

ERNA and eDNA were co-extracted using the AllPrep DNA/RNA Mini Kit (Qiagen, Hilden, Germany). The cryopreserved Sterivex cartridges were thawed at 4 °C and the RNAlater removed by centrifugation in a 50 mL tube at 8000 g for 1 min before introducing 1.5 mL of lysis buffer (Buffer RLT Plus) through the inlet. Cartridges were then incubated under mild rotation at 60 °C for 60 min and the lysis buffer was collected by centrifuging in a 50 mL tube and transferring to an AllPrep DNA spin column. The subsequent steps were conducted using an elution volume of 50 μL according to the manufacturer’s protocol. DNA samples were purified using the NucleoSpin DNA Clean-up XS kit (Macherey-Nagel, Düren, Germany). Before cDNA synthesis, RNA samples were subjected to two consecutive DNase treatments with an rDNase set and the NucleoSpin RNA Clean-up XS kit (MACHEREY-NAGEL, Düren, Germany). cDNA was synthesized from single-stranded RNA using the PrimeScript II 1st Strand cDNA Synthesis Kit (TaKaRa Bio Inc., Shiga, Japan) according to the manufacturer’s protocol. DNA and RNA were extracted simultaneously from the distilled water to assess cross-contamination during sample filtration, co-extraction, and cDNA synthesis. All extracted DNA/RNA and cDNA samples were immediately stored at −80 °C.

### 2.5. Amplicon library preparation and sequencing for eDNA/eRNA metabarcoding analysis

Amplicon libraries of partial 16S rRNA genes were obtained by PCR amplification using Takara Ex Taq Hot Start Version (TaKaRa Bio Inc., Shiga, Japan) and the MtInsects-16S primer set for insects in the mtDNA 16S rRNA region (Table S2)^34,35^. The first PCR was carried out in 10 μL of the reaction mixture with 6.1 μL of sterile distilled H_2_O, 1 μL of 10 × PCR Buffer, 0.8 μL of dNTP mixture, 0.5 μL of each MtInsects-16S primer (10 μM), 0.1 μL of Takara EX Taq HS, and 1 μL of the DNA or cDNA template. Thermal cycling was performed at 94 °C for 1 min followed by 35 cycles at 94 °C for 1 min, 50 °C for 30 s, 72 °C for 30 s, and 72 °C for 3 min. PCR amplification was performed eight times for each DNA and cDNA sample. Purification was performed using the AMPure XP System (Beckman Coulter, Brea, CA, USA). The second PCR was then conducted using 10 µL of the reaction mixture with 3.1 μL of sterile distilled H2O, 1 μL of 10 × Ex Taq Buffer, 0.8 μL of dNTP mixture, 2 μL of each primer (2.5 μM) including the Nextseq adaptor sequences and 8-bp index sequences at each amplicon end, and 1 μL of the DNA or cDNA template. Thermal cycling was performed at 94 °C for 2 min, followed by 10–12 cycles at 94 °C for 30 s, 60 °C for 30 s, 72 °C for 30 s, and 72 °C for 5 min. The PCR products were purified using the AMPure XP System. PCR controls were used to assess cross-contamination in both rounds. Equal amounts of purified amplicons were then mixed, and libraries for eRNA and eDNA produced as 2 × 300 bp paired-end sequencing on a NextSeq 2000 instrument using NextSeq 2000 P1 Reagents (600 cycles) (Illumina, San Diego, CA, USA) according to the manufacturer’s instructions.

### 2.6. Bioinformatics analysis of high-throughput sequencing data

Bioinformatics analyses were performed using the Quantitative Insights into Microbial Ecology (QIIME2 v. 2021.4) pipeline. Primer sequences were removed using the Qiime2 cutadapt plugin (qiime cutadapt trim-paired). The divisive amplicon denoise algorithm 2 (DADA2) denoising pipeline from QIIME2 was used for denoising the sequence reads, chimeric sequence, singleton, doubleton, and tripleton removal, and forward and reverse paired-end read merging to generate feature tables of the amplicon sequence variants (ASVs). A BLAST search was performed on all processed sequences using local BLAST 2.15.0+ against the reference DNA database, which is integrated with the database for insects created by Kanagawa Prefecture English version and GenBank^35^. Details of the procedure for creating this DNA database are described on GitHub^36^. The top-scoring species name was assigned for each ASV with a >97% sequence identity (as the product of the percentage of identical matches (pident) and query coverage per subject (qcovs)) to the database. ASVs below an identity threshold of 97% were classified as “unknown” and eRNA and eDNA with read numbers of < 10 bp were discarded. Target species in the benthic macroinvertebrate survey in the Ministry of Land Infrastructure and Transport-River Environmental Database (MLIT-RE database) were extracted from the organism list detected by eNA metabarcoding and the resulting ASV and taxonomy tables of those species used for further analyses^37^. The species name output from the BLAST search was manually confirmed using information from the literature and corrected to the most recent scientific name. Habitats (aquatic, terrestrial, or both) were also classified using information from the literature (Tables S3 and S4). Literatures used for confirmation of species names and habitat classifications were summarized in Text S2. All raw sequences were deposited in the NCBI Sequence Read Archive under accession number XX.

### 2.7. Traditional field survey (TFS)

Qualitative and quantitative ecological surveys of arthropods were conducted in December according to the manual from the National Census of River Environments^38^. Samples were collected by kick sampling or wash sampling using dip nets in each habitat area (e.g., riffles and pools) for the qualitative study, while quantitative studies were conducted using 25 cm × 25 cm (0.5 mm mesh) Surber samplers in the rapids at each sampling site. The results were combined to create a list of the presence/absence and abundance of species observed at each sampling site.

### 2.8. Calculation of biodiversity indicators based on species detected by TFS and eNA metabarcoding analysis and estimation of the safe concentration (SC) of LAS

ENA metabarcoding analysis and TFS datasets were used to calculate the total families, total taxa, average score per taxon (ASPT), and percentage of Ephemeroptera, Plecoptera, and Trichoptera (EPT) taxa (%EPT) to evaluate changes in the community structures at the different sampling sites. The ASPT score was calculated according to the Manual for Water Quality Assessment of Aquatic organisms in Japan^39^, while %EPT was calculated based on the proportion of species classified as Ephemeroptera, Plecoptera, and Trichoptera divided by total number of species detected at each sampling site^40,41^. The SC for LAS was estimated by comparing the indices at the different sampling sites with different LAS concentrations.

### 2.9. Statistical analyses

Statistical analyses were performed using Origin Pro 2021b (OriginLab, Northampton, MA, USA) or R version 4.3.1 (R Core Team, Vienna, Austria) with the minimum significance level set at *p* = 0.05. The normality of each dataset was assessed using the Shapiro-Wikl test. Water quality and biological indices (total families, total taxa, ASPT, and %EPT) for the different sampling sites were compared using one-way analysis of variance (ANOVA) (for parametric and homoscedasticity cases), Welch’s one-way ANOVA (for parametric and heteroscedasticity cases), or the Kruskal-Wallis test (for non-parametric cases). Post-hoc analysis was conducted following statistical analysis using Tukey’s test, the Games-Howell test, or Dunn’s test. A Venn diagram was generated to compare the number of species detected by eRNA/eDNA metabarcoding and TFS. Some species detected by eRNA and eDNA that were difficult to identify at the species level under TFS sampling were renamed to the family or genus level, as illustrated in the Venn diagrams (Table S5). To examine the performance of eRNA/eDNA metabarcoding analysis, sensitivity and positive predictivity were calculated using the following equation^24,25^.

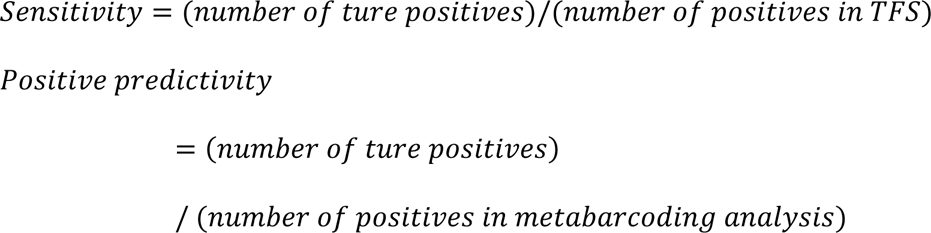

Heatmap and cluster analyses based on the Jaccard distances were then constructed using the R package pheatmap (function: “pheatmap”) to visualize the presence/absence of the 179 species detected by TFS in the eRNA/eDNA datasets (Table S6). Scatter plots of the eRNA and eDNA read numbers were generated for all detected species (Tables S3 and S4). The average read numbers at the three sampling sites were then calculated and 1 added to enable the use of a logarithmic scale. To discuss the ability of eNA to distinguish false-positive species, terrestrial species were defined as potential false-positives that were unlikely to inhabit the aquatic environment at the sampling sites. The number of species and false-positive rates were compared using a paired-sample *t*-test because the variance was homogeneous. The presence/absence of datasets detected by eRNA and eDNA analyses were used in non-metric dimensional scaling (nMDS) and redundancy analysis (RDA), excluding terrestrial species. NMDS analysis was performed based on the Jaccard distances to examine differences in the species composition for the eRNA and eDNA between the sampling sites using the R package vegan (function: “metaMDS”). The significance of the results was then tested with pairwise multilevel comparisons to evaluate where differences between benthic communities existed using the R package pairwiseAdonis (functon: “pairwise.adonis”). RDA was used to examine the effects of water quality parameters on the benthic communities at the different sampling sites. Water quality parameters were log-transformed to normalize the variable distributions when necessary. To avoid problems with multicollinearity, RDAs were conducted using a forward selection procedure in the R package vegan (function: “ordistep” and “rda”), while variance inflation factors (VIF) were calculated on the variables from the RDA using the R package vegan (function: “vif.cca”) to identify redundancy in any of the selected variables^42^. The significance between benthic communities and water quality parameters was tested using the R package stats (function: “anova”).

## 3. RESULTS AND DISCUSSION

### 3.1. Comparison of water quality at different sampling sites

Hazard assessment using field surveys such as TFS and eNA analysis is associated with issues in data variability and difficulty in interpreting results due to problems with controlling confounding factors; however, it is possible to minimize these effects by sampling at contaminated and non-contaminated sites with similar physicochemical conditions^43,44^. The water quality parameters at the different sampling sites were thus compared to evaluate any similarities in the physicochemical conditions. The Ministry of the Environment in Japan has established environmental quality standards for conservation of the living environment^45^, with the water quality indicators used to characterize rivers; pH, BOD, DO, SS, and the number of *E. coli*, used to classify environmental quality standards from the cleanest Class AA to the dirtiest Class E. The water quality at the sampling sites was thus evaluated based on these criteria (Figure 2). All sites were classified Class A based on the average pH (upstream: 8.1, outfall: 8.2, downstream: 8.2), BOD (upstream: 0.25, outfall: 1.25, downstream: 0.29 mg/L), DO (upstream: 13.6, outfall: 12.8, downstream:13.6 mg/L), SS (upstream: 0.50, outfall: 0.75, downstream: 0.50 mg/L), and number of *E. coli* (upstream: 0.92, outfall: 0.67, downstream: 49 CFU/100mL), indicating good water quality, However, BOD tended to be relatively higher at the outfall site. The 95th percentiles for LAS concentration were 0.21, 47, and 11 µg/L at the upstream, outfall, and downstream sites, respectively, demonstrating significantly higher LAS concentrations at the outfall and downstream sites as compared to the upstream site. TOC was also significantly higher at the outfall and downstream sites, with significant differences observed between the upstream and outfall sites (Figure S1). Based on the PRTR information, only LAS is contained in the effluent from the plant. Therefore, the increases in LAS concentration, BOD, and TOC at the outfall and downstream sites can be attributed to LAS in the plant effluent. Among the water quality parameters, the number of *E. coli* tended to be significantly higher at the downstream site; however, this may be due its presence in compost from fields in the vicinity of the downstream site. However, its effect on aquatic organisms was considered low because the average number of *E. coli* downstream was within Class A levels. The results indicate that the river water in the sampling sites was clean enough to be classified Class A, with the LAS concentration relatively high downstream of the plant effluent outfall, meaning that the selected site is suitable for investigating the effect of LAS as a single contaminant.

**Figure 2.**
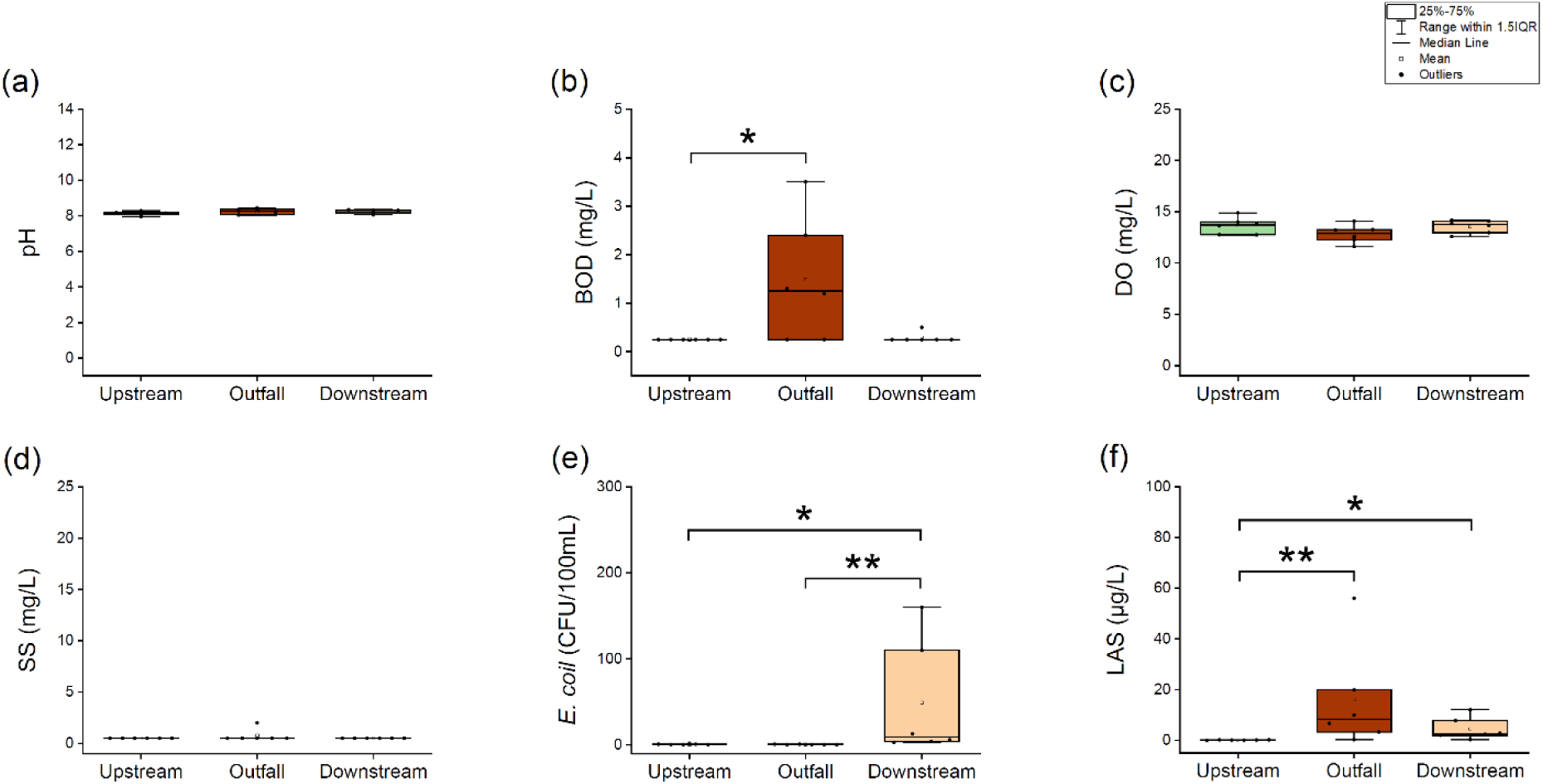
Comparison of water quality at different sampling sites: (a) pH, (b) BOD, (c) DO, (d) SS, (e) the number of E. coli, and (f) LAS concentration. Differences were analyzed using one-way analysis of variance or the Kruskal-wallis test. Asterisks indicate factors that are statistically significant after applying multiple comparison adjustments for Turkey’s test or Dunn’s test, respectively (*: *p*<0.05, **: *p*<0.01).

### 3.2. Differences in species detected using eRNA/eDNA analyses vs TFS

The obtained reads were clustered into 422 ASVs for eRNA and 483 ASVs for eDNA sequences (Tables S3 and 4). All negative controls (filtration, extraction, reverse transcription, and PCR) were merged and sequenced for eRNA and eDNA, with no species detected. Difficulties in identification at the species level when sampled by TFS meant that some species detected by eNA were renamed at the family or genus level for comparison (Table S5).

Differences in the species detected at the three sampling sites when using eNA metabarcoding and TFS were illustrated in a Venn diagram (Figure. 3a–d, Table S5). ERNA and eDNA analyses detected 372 and 430 species, respectively, while TFS detected 179 species (Figure 3a). The number of species identified using eRNA and eDNA was thus 2.1 and 2.4 times that identified using TFS, respectively, which is consistent with previous studies^35^. Although eRNA disappears more quickly than eDNA, eNA is thought to remain in water for long periods, and because eNA metabarcoding analysis can detect even low concentrations of eNA^46,47^, it may have the ability to capture historical data about organisms or detect species with low abundance that could not be sampled by TFS. The sensitivity of eRNA and eDNA analyses was 56%, and the positive predictive value of eRNA (27%) was slightly higher than that of eDNA (23%) at all three sampling sites. Interestingly, a lower number of eRNA species was detected at the outfall and downstream sites as compared to eDNA (Figure 3c and d), whereas the number of eRNA species detected was higher upstream (Figure 3b). These results indicate that RNA may be more abundant than DNA in the nucleic acids released by living organisms in aquatic environments. ENA primarily disappears from the environment through biodegradation by microorganisms and photodegradation^48,49^, and because RNA is more degradable than DNA^50,51^, the number of species detected by eRNA is generally less than that detected by eDNA. However, more species were detected by eRNA than eDNA at the upstream site. This may be because of a relatively small number of microorganisms in the upstream site, as evidenced by lower BOD and TOC, with the lower amounts of organic substances limiting the energy available for microorganisms. Assuming that more RNA is released by organisms than DNA, eRNA may have a higher detection ability than eDNA in an aquatic environment with low organic substances and few microorganisms because biodegradation is less likely in a clean aquatic environment. To verify this hypothesis, detailed studies on the mechanisms of nucleic acid release from organisms are required.

**Figure 3.**
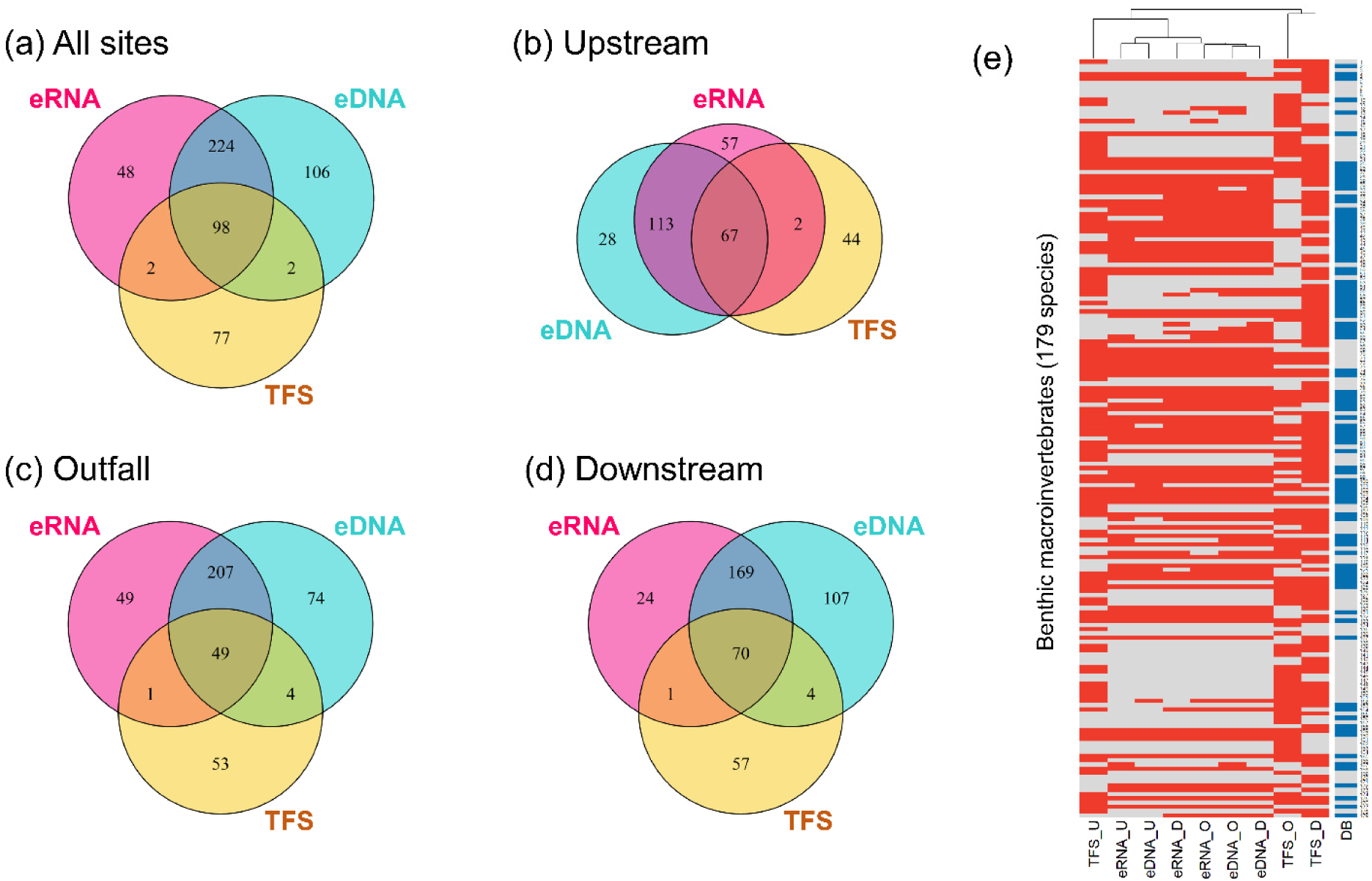
Comparison of species based on eDNA and eRNA metabarcoding analyses and TFS. Venn diagrams indicate the number of species detected at (a) all sites, (b) the upstream site, (c) the outfall site, and (d) the downstream site. (e) Heatmap and cluster analyses of benthic compositions detected by TFS, eRNA, and eDNA upstream, outfall, and downstream. Rows correspond to benthic macroinvertebrates detected by TFS (179 species) and columns to species detected by TFS, eRNA, or eDNA (U: upstream, O: outfall, D: downstream). Heatmap colors indicate the presence/absence of species (red: presence, gray: absence). The distance matrix was calculated using Jaccard distances. DB indicates whether a species is registered in the reference DNA database (blue: registered, gray: unregistered).

Heatmap analysis of the TFS and eRNA/eDNA metabarcoding, based on the presence/absence datasets at each sampling site, revealed distinct compositional differences between the TFS and eRNA/DNA datasets (Figure 3e, Table S6). The biological communities detected by eRNA/eDNA analysis clustered relatively close to the community observed by TFS at the upstream site. Previous studies have found that eDNA remains in the aquatic environment for long periods and can be transferred at least 2–3 km upstream to downstream^52–54,26^, meaning that eNA from organisms living in the upstream site may be detected at the outfall and downstream sites, approximately 400 m and 2000 m away, respectively. Of the 12 species collected only upstream by TFS and registered in the reference database, 11 and 11 were detected at both the outfall and downstream sites by eRNA analysis, respectively, and 10 and 11 by eDNA analysis (Figure 3e, Table S6). It is therefore possible that the communities detected downstream by eNA metabarcoding were similar to those collected upstream by TFS because eNA from organisms upstream was transported downstream. When comparing the eRNA and eDNA communities, those detected using eRNA at the upstream, outfall, and downstream sites clustered separately, whereas those detected using eDNA at the outfall and downstream sites clustered in the same group. This suggests that eDNA may be difficult to capture changes in the benthic community alongside changes in LAS concentration. The community detected by TFS at the outfall and downstream sites also clustered relatively far from that detected by eNA metabarcoding, suggesting that several of the species sampled by TFS were not detected by eNA metabarcoding. To confirm these details, we checked whether the 77 species detected by TFS, but not by eNA, were registered in the reference DNA database, and found that only 12 species were registered and 65 species (84%) were not. Limiting to the species detected by TFS and registered in the reference DNA database (114 species), the sensitivity of eRNA and eDNA was 88%, suggesting that eNA analysis is highly sensitive for detecting species registered in the reference database. Therefore, the failure to detect TFS species by eNA analysis could be because they are not registered in the reference DNA database, and expanding the sequence database may improve the detection sensitivity of eNA metabarcoding^25^. In addition, of the 12 species that were registered in the reference DNA database but could not be detected by eRNA/DNA, eight (*Nais communis*, *Epiophlebia superstes*, *Planaeschna milnei milnei*, *Anisogomphus maacki*, *Hexatoma* sp., *Microtendipes* sp., *Stictochironomus* sp., and *Dixa nipponica*) are found in river sand or mud. Because only surface water was sampled in this study, it is possible that organisms living in the sediment were not detected. Hence, the development of a new approach for collecting eDNA and eRNA from benthic organisms is necessary to improve the detection sensitivity of eNA analyses. For instance, previous studies have reported that bulk samples are a superior source of benthos eNA compared with water samples^55^. For a comprehensive biodiversity assessment, it may be more effective to conduct both water and bulk sample analyses to ensure that eNA are captured from all target organisms.

### 3.3. Comparison of the false positive rates in eRNA and eDNA analysis

Comparison of the relative abundances of eDNA and eRNA reads at the sampling sites shows that the eDNA read numbers correlate significantly with eRNA (R=0.82, *p*<0.01) (Figure 4a, Tables S3 and 4). This trend is also consistent in previous studies^24^, and indicates sufficient eRNA for metabarcoding analysis. In total, 366 species were detected using both eRNA and eDNA, and 56 and 117 species were detected using only RNA and DNA, respectively (Figure 4a). Of these, 104, 28, and 85 terrestrial species were obtained as false positives, respectively, indicating that eDNA analysis detected a relatively large number of terrestrial species. The number of species detected in each eRNA/eDNA sampling event (3 sites × 6 times) was not significantly different in terms of eRNA and eDNA (Figure 4b); however, the detection rate of terrestrial species, as false positives, was significantly higher for eDNA (range: 5–34 %) than eRNA (range: 2–25 %) (Figure 4c). EDNA lasts for relatively long periods; thus, there may be an abundance of eDNA derived from dead terrestrial organisms or living terrestrial organisms on riverside soils or tree leaves, explaining the detection of terrestrial species in the water samples, while the relative instability of eRNA reflects short-term benthic communities and reduces false positives. As described in the previous section, eNA can be used to detect resident species with high sensitivity, and eRNA is less likely to detect false-positives as compared to eDNA; thus, eRNA analysis is more useful for biological monitoring that focuses solely on aquatic species.

**Figure 4.**
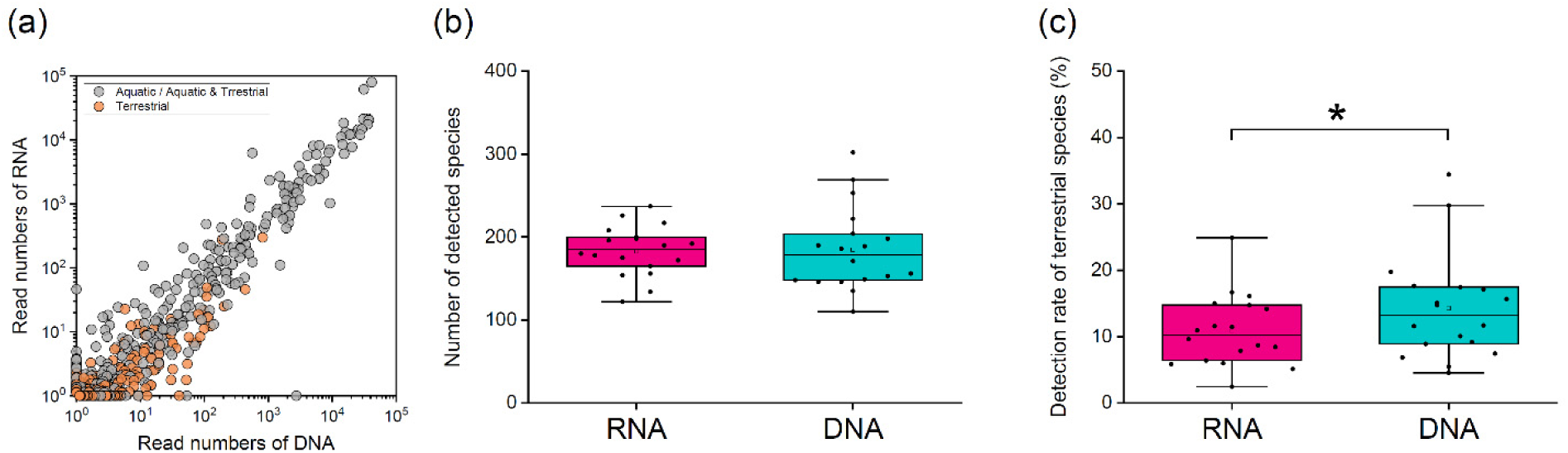
Comparison of eRNA and eDNA in reducing false positives. (a) Scatterplot of average numbers of eRNA and eDNA reads obtained for benthic macroinvertebrates at all sites. Box plots show (b) number of benthic macroinvertebrates and (c) detection rate of terrestrial species per sampling day (n=18). Differences were analyzed using the paired-sample t-test (*: *p*<0.05).

### 3.4. Beta diversity of benthic community and related water quality parameters

Beta diversity is an important indicator of the relationships between benthic communities and environmental factors^56,57^. In this study, the beta diversity was analyzed based on presence/absence datasets, excluding terrestrial species, with nMDS used to confirm the similarity between the benthic communities at the different sampling sites. The nMDS analysis of eRNA revealed remarkable site-to-site variation in the benthic communities (Figure 5a). Plots of the upstream and outfall sites are concentrated in a negative direction on the NMDS1 axis, whereas those describing the downstream site are concentrated in the positive direction. Additionally, plots of the upstream and outfall sites are concentrated in the negative and positive direction on the NMDS2 axis, respectively, whereas those describing the downstream site are concentrated between the upstream and outfall sites along the NMDS2 axis. The PERMANOVA results confirmed significant differences in the benthic communities between the upstream and outfall sites (R^2^ = 0.253, *p* = 0.006; Table S7), and between upstream and downstream sites (R^2^ = 0.364, *p* = 0.015; Table S7). Water was clean at all sampling sites; however, LAS was detected at relatively high concentrations downstream of the chemical plant effluent outfall. Therefore, the benthic communities in the outfall and downstream sites may be affected by LAS. The nMDS analysis of eDNA also showed significant differences between the upstream and outfall/downstream sites, similar to eRNA, although variation was observed at the outfall and downstream sites in each plot (Figure 5b, Table S8). These results suggest that eNA analysis has the potential to detect community variations at different sampling sites, with differences in the communities at the upstream and outfall sites distinguished at a distance of 400 m in this study. Surprisingly, the *p*-values indicating the differences in the benthic communities at the upstream and outfall sites were smaller for eRNA (*p* = 0.006; Table S7) than for eDNA (*p* = 0.030; Table S8). The stability of eDNA may have led to the detection of false-positive species and upstream-flowing eDNA, which may have prevented clear separation between the communities in the sampling sites in the nMDS analysis, verifying that eRNA is more sensitive than eDNA for detecting community differences.

**Figure 5.**
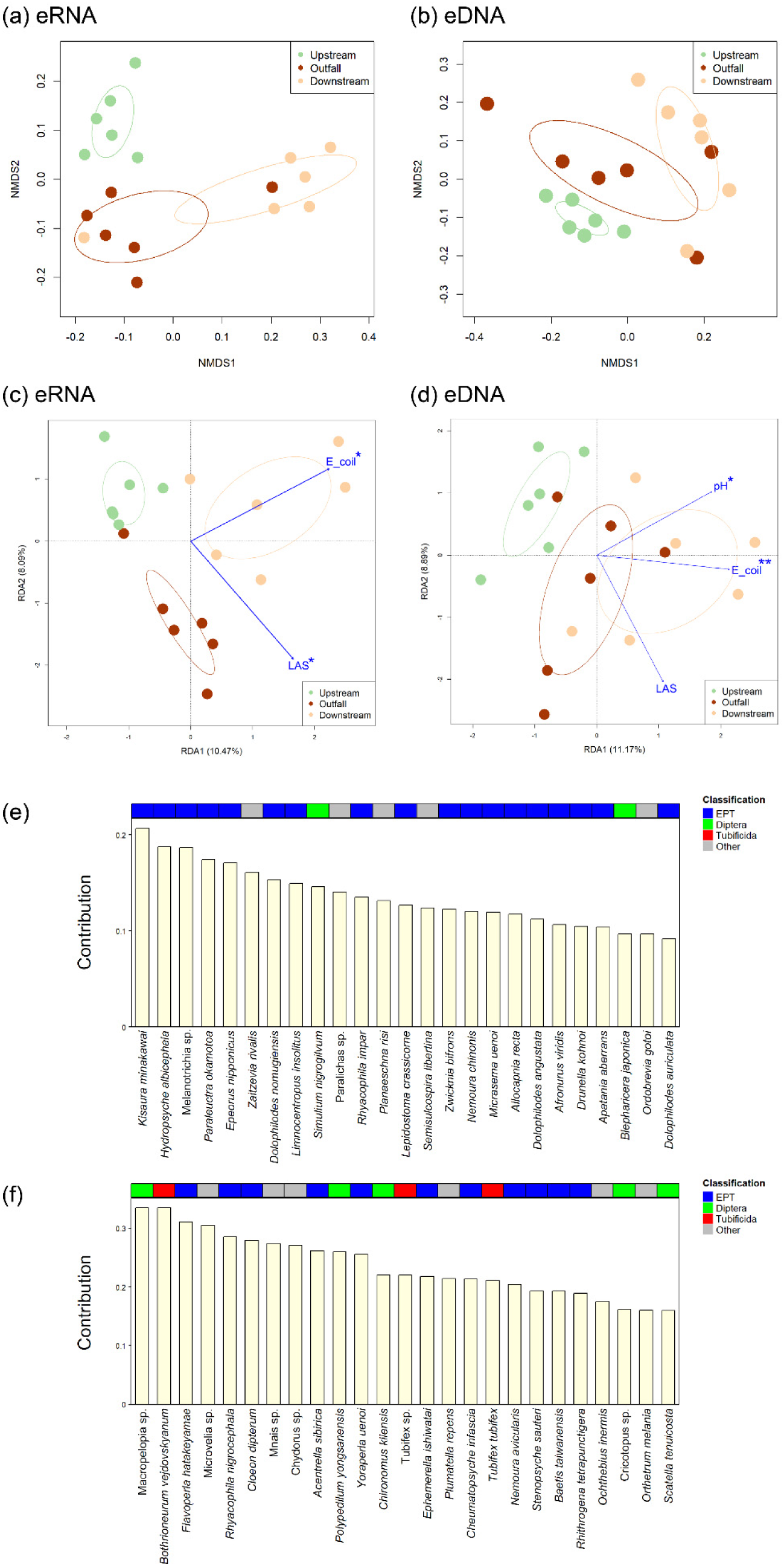
Comparison of beta diversity for benthic community and related water quality parameters at different sampling sites, with non-metric multidimensional scaling ordination (nMDS) of the benthic communities detected by (a) eRNA and (b) eDNA analysis at each sampling site. Analysis demonstrated high credibility for the ordination results (eRNA: stress = 0.11, eDNA: stress=0.12). Redundancy analysis (RDA) of benthic communities, species, and environmental factors in the sampling sites detected by (c) eRNA and (d) eDNA analysis. Direction of blue arrow indicates positive correlation between the environmental factors obtained by forward selection and the benthic communities. Asterisks indicate statistically significant factors (*: *p*<0.05, **: *p*<0.01). Green plots in both nMDS and RDA indicate samples from the upstream site, dark orange plots indicate samples from the outfall site, and light orange plots indicate samples fromdownstream site. Ellipses indicate standard errors with 95% similarity for each sampling site. Bar chart shows top 25 contributing species that correlate (e) negatively and (f) positively with LAS and their contribution based on species scores in the RDA (Table S11 and S12). Blue annotations indicate Ephemeroptera, Plecoptera, and Trichoptera (EPT) species while green annotations indicate Diptera and red annotations Tubificidae.

Changes in the biological communities are due to multiple environmental factors. Previous studies have found that pH, DO, and organic pollution are the main environmental factors affecting benthic communities^58^. The nMDS results suggested that communities in the outfall and downstream sites may have been affected by LAS because of relatively high LAS concentrations. RDA was performed to understand the reasons for the differences in benthic communities at the sampling sites by analyzing the relationship between the communities and water quality parameters (Figure 5c and d). The forward selection procedure indicated that, of the six water quality parameters (pH, BOD, DO, SS, number of *E. coli*, and LAS concentration), the LAS concentration and number of *E. coli* were the main factors influencing the benthic communities detected by eRNA analysis (Figure 5c). Collinearity test showed VIFs of 1.06 and 1.06 for the LAS concentration and the number of *E. coli*, respectively, indicating no collinearity. The RDA results for the eRNA analysis showed that the first two axes explained 18.56% (adjusted R^2^ = 7.70%) of the total variance (RDA1 = 10.47%, RDA2 = 8.09%). The benthic community was significantly linked to the LAS concentration (*F* = 1.58, df = 1, *p* = 0.032; Table S9) and number of *E. coli* (*F* = 1.83, df = 1, *p* = 0.014; Table S9). The number of *E. coli* on the first axis wass associated with the benthic communities, while the second axis indicated the relationship between LAS concentration and the benthic communities. The results show positive correlation between the number of *E. coli* and the benthic community at the downstream site, with *Tinearia alternata*, *Mataeopsephus japonicus*, *Potthastia sp.*, *Simulium quinquestriatum*, and *Sympecma paedisca* positively correlated with the number of *E. coli* (Table S10). Of the top five contributors, *T. alternata* and *M. japonicus*, of the families Psychodidae and Psephenidae, respectively, are indicator species that inhabit slightly dirty aquatic environments, while *Potthastia sp.* and *S. quinquestriatum*, of the families Chironomidae and Simuliidae, respectively, are indicator species that inhabit dirty aquatic environment^39,59^. The number of *E. coli* at the downstream site was significantly higher than that at the upstream/outfall sites (Figure 2e). If the number of *E. coli* exerted a great influence on the benthic community, significant differences should be observed not only between the upstream and downstream sites, but also between the outfall and downstream sites; however, no such differences were observed (Figure 5a, Table S7). Additionally, the number of *E. coli* fell within that allowed for Class A water quality (<300 CFU/100 mL); thus, the number of *E. coli* might not markedly affect biological communities^45^. On the other hand, the results of RDA and nMDS showed that LAS was significantly correlated with benthic communities (Figure 5c), with the communities differing significantly in the upstream and outfall/downstream sites (Figure 5a, Table S7). The LAS concentration was positively correlated with the community at the outfall site and negatively correlated with the community upstream. To understand the characteristics of species that correlated with LAS, those plotted in the second and fourth quadrants of the RDA results were identified. Fifteen (75%) of the top 25 species plotted in the second quadrant, which were negatively correlated with LAS, were EPT species, which are commonly used as indicators of environmental river health because of their high sensitivity to deteriorated water quality^30,60^ (Figure 5e, Table S11). Two (0.1%) and zero (0%) of the species were Diptera and Tubificidae, respectively, which indicate dirty environments^59,61,62^. Species positively correlated with LAS were observed in the fourth quadrant, with 11 (44%) of the top 25 composed of EPT species (Figure 5f, Table S12); however, 5 (20%) and 3 (12%) Diptera and Tubificidae, respectively, were observed, which is more than that in the second quadrant. These results suggest that organic pollution caused by LAS in plant effluent may render it difficult to detect EPT species while facilitating the detection of Diptera and Tubificidae; therefore, LAS may be affecting sensitive species such as EPT in association with the water quality downstream of the plant, particularly at the outfall site where the highest concentrations of LAS were detected. *Macropelopia sp.*, *Bothrioneurum vejdovskyanum*, *Flavoperla hatakeyamae*, *Microvelia sp.*, and *Rhyacophila nigrocephala* were positively correlated with LAS concentration (Figure 5f, Table S12), whereas negative correlations were observed with *Kisaura minakawai*, *Hydropsyche albicephala*, *Melanotrichia sp.*, *Paraleuctra okamotoa*, and *Epeorus nipponicus* (Figure 5e, Table S11). Species that are highly correlated with LAS may therefore act as indicator species to visualize the effects of LAS in real aquatic environments.

For eDNA metabarcoding, RDA explained 25.58% of the total variance (with an adjusted variation of 9.64%). The number of *E. coli*, LAS concentration, and pH were the main water-quality parameters selected for use with the forward selection procedure (Figure 5d, Table S13). The VIFs for the number of *E. coli*, LAS concentration, and pH were found to be 1.15, 1.08, and 1.13, respectively, indicating no collinearity. The number of *E. coli* and pH were associated with the community structures of the sampling sites on the first axis, while the second axis indicated the relationship between the LAS concentration and the benthic community. The number of *E. coli* and LAS concentration tended to be positively correlated with the benthic communities at the downstream and outfall sites, respectively; however, the communities at each site did not plot separately (Figure 5d). This result was also supported by nMDS for eDNA. The number of *E. coli* (*F* = 1.81, df = 1, *p* = 0.008) and pH (*F* = 1.48, df = 1, *p* = 0.044) were key variables in determining the characteristics of the benthic community (Table S13). Surprisingly, however, no significant correlation was observed between the LAS concentration and benthic communities (*F* = 1.51, df = 1, *p* = 0.056), whereas a significant correlation was observed in the RDA analysis of eRNA. These results suggest that eRNA can accurately capture changes in benthic communities along with changes in environmental factors because eRNA is unstable compared to eDNA and reflects short-term benthic communities.

### 3.5. Ecological effect assessment of LAS based on analysis of biological communities using TFS and eRNA/eDNA, and estimation of the SC of LAS

The most common method for the hazard assessment of chemicals is the AF method, in which the PNEC is calculated by extrapolating laboratory-measured toxicity data for individual species to real ecosystems. However, few studies verify whether chemicals have a negative impact on real ecosystems at the PNEC^14,63^. Because the selected river is a site at which the effect of LAS alone on the benthic macroinvertebrates in the environment may be evaluated, we estimated the SC for LAS based on changes in the total families, total taxa, ASPT score, and %EPT, which were calculated from a wide variety of benthic macroinvertebrates at sites with different LAS concentrations, and compared them with the PNEC to verify the validity of the AF method (Figure 6).

**Figure 6.**
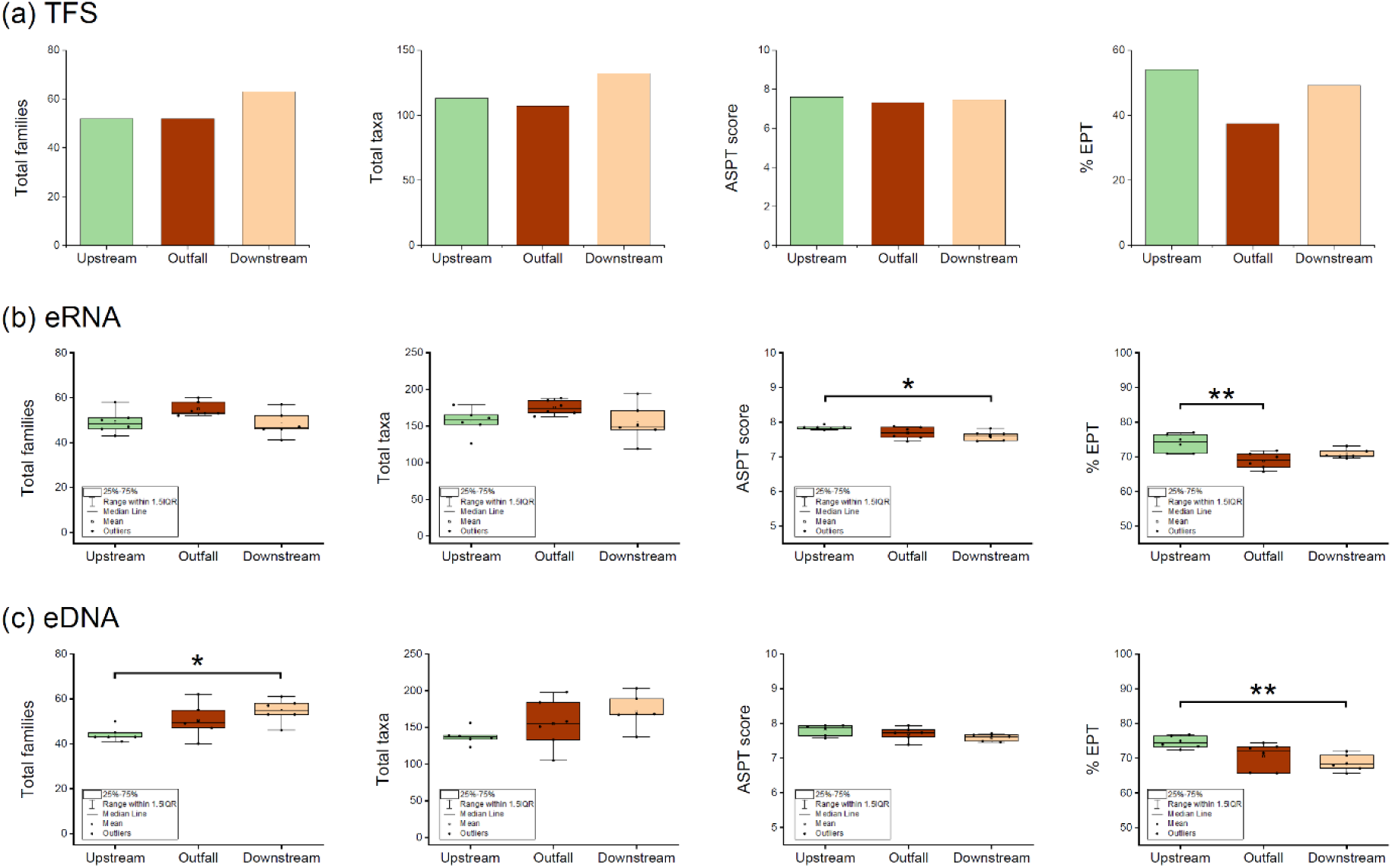
Comparison of various indicators at different sampling sites. Bar plots show total families, total taxa, ASPT score, and %EPT based on the species detected by (a) TFS, while box plots indicate total families, total taxa, ASPT score, and %EPT based on species detected by (b) eRNA and (c) eDNA analysis. Differences in indicators were analyzed by one-way analysis of variance (ANOVA) or Welch’s one-way ANOVA. Asterisks indicate factors that are statistically significant after applying the multiple comparison adjustments for Turkey’s test or Games-Howell test, respectively. (*: *p*<0.05, **: *p*<0.01).

The total number of families and taxa detected by TFS changed little at the upstream and outfall sites; however, an increase was observed downstream (Figure 6a). As mentioned in Text S3, the number of detections may increase because the downstream site is a suitable environment for species that inhabit both clean and slightly dirty environments. The ASPT scores used as water quality indices ranged from 7.3 to 7.6, and the differences observed between the different sampling sites were unremarkable, indicating good water quality at each site. However, the %EPT was relatively low at the outfall site and tended to recover at the downstream site.

The total families (average value: upstream: 49, outfall: 55, downstream: 48) and taxa (average value: upstream: 156, outfall: 175, downstream: 155) were calculated based on the species detected by eRNA analysis, and these indices did not differ significantly at the different sampling sites (Figure 6b). Although significant differences were observed for the ASPT scores at the upstream and downstream sites, the average values (upstream: 7.8, outfall: 7.7, downstream: 7.6) indicated clean water quality and were similar to those obtained by TFS (Figure 6b). Notably, no significant differences were observed in these indices between the upstream and outfall sites. Because the total families, total taxa, and ASPT scores target a wide variety of organisms that inhabit various environments (i,e., indices that can measure the impact on the entire ecosystem), these indices should change significantly between the upstream and outfall sites if LAS is impacting the entire ecosystem in the study site. Therefore, the LAS concentrations detected in this study were considered to have little effect on the entire ecosystem. However, the %EPT at the outfall site (average value: 69) was significantly decreased as compared to that upstream (average value: 74), and a recovery trend was observed downstream (average value: 71). This trend corresponds to the TFS results. Ephemeroptera, Plecoptera, and Trichoptera are sensitive to water pollution and are often used as indicators of water quality^60^. These results suggest that LAS may have affected some of the water pollution-sensitive species. However, this effect might not be fatal because no significant differences were observed in the %EPT at sites located 1650 m downstream. Comparison of these indices based on the eDNA dataset indicated no significant differences were observed in the ASPT scores between the sampling sites (average value Upstream: 7.8, Outfall: 7.7; downstream, 7.6). However, an increasing trend in the total number of families (average value: upstream: 44, outfall: 51, downstream: 55) and taxa (average value: upstream: 138, outfall: 155, downstream: 172) was observed downstream (Figure 6c). Additionally, significantly reduced %EPT was observed downstream as compared to upstream, and a recovery trend for %EPT was not observed. The results of nMDS and RDA suggest that eDNA analysis reflects the long-term benthic composition and detects eDNA derived from macroinvertebrates upstream because of its stability, which may have led to the increase in the number of total families and taxa and decrease in %EPT at the downstream site.

The LAS concentration measurement indicated 95th percentiles of 0.21, 47, and 11 µg/L for the upstream, outfall, and downstream sites, respectively. The estimated SC of >47 µg/L for LAS at the outfall site, which was obtained from the eRNA datasets and is consistent with TFS, indicated no effects on the total families, taxa, or ASPT scores. In previous studies, the calculation of PNEC using the AF method (PNEC_AF_) for LAS reported as 2.8 µg/L^64^, which is lower than the SC estimated in this study. One study investigating C12-LAS over a 56-day experimental stream mesocosm study reported a NOEC of 268 µg/L^65^ and the mesocosm PNEC of 268 µg/L (NOEC/AF, AF=1)^65,66^. The SC calculated in this study was thus closer to the mesocosm PNEC than to the PNEC_AF_. Although it is difficult to determine which PNEC values should be used as the standard concentration for biodiversity conservation, the AF method may result in large uncertainties in the assessment results because the impacts on ecosystems are predicted by extrapolation from the effects of a single species^12–15^, leading to conservative assessment of the effects of chemicals on ecosystems. On the other hand, the SC to protect the water pollution-sensitive species such as EPT was estimated at 11–47 µg/L because the %EPT was significantly lower at the outfall site than the upstream site. Protecting sensitive and endangered species is important for biodiversity conservation, and it is essential to establish individual water quality standards for rivers that are inhabited by such species to prevent loss. ERNA analysis may be useful in setting this standard, because it is highly sensitive in capturing the responses of biological communities to changes in the water quality.

These results of community-level hazard assessment using eRNA could provide important knowledge that complements the assessment results using the AF method. To properly manage chemical substances, the results of community-level hazard assessments should be actively incorporated as a priority and discussed to verify the validity of current risk assessment methods and environmental standard^67^.

As described above, eNA helps to understand the effects of chemicals and may contribute to the derivation of appropriate criteria for the development of water quality standards. In particular, eRNA analysis is sensitive in capturing the responses of biological communities and may therefore be useful in visualizing the relationship between ecosystems and anthropogenic chemical contamination, and in assessing ecosystem restoration through reducing contamination. ENA metabarcoding could become a more powerful biomonitoring tool, provided the following limitations are addressed. First, the sequencing and registration of species not detected by eNA metabarcoding. Second, the development of a new approach for collecting eNA from benthic organisms is necessary. Third, transport distance of eNA in nature river should be understood. If these limitations can be overcome, the detection sensitivity and positive predictivity may be improved. Additionally, this is the first study to assess the hazards of chemicals based on the responses of benthic communities via eNA metabarcoding in real aquatic environments. The issues in hazard assessment using the AF method will become clearer by case studies using eNA analysis for other chemicals.

## ASSOCIATED CONTENT

The Supporting Information is available free of charge on the ACS Publications website.

Measurements of LAS concentrations and the literature used for the confirmation of species names and habitat classification are presented in the Supplementary Text.

Comparison of TOC between sampling sites (Figure S1); Cluster analysis result of benthic communities based on the abundance data of TFS (Figure S2); Dates of field survey and water quality measurements obtained (Table S1); MtInsects-16S primer pairs used for metabarcoding (Table S2); PERMANOVA analysis of benthic communities between sampling sites detected by eRNA (Table S7) and eDNA (Table S8) in nMDS; Results of significant test of correlation between water quality parameters and benthic communities detected by eRNA (Table S9) and eDNA (Table S13) (PDF)

Read numbers of species detected by eRNA (Table S3) and eDNA (Table S4) analyses; Presence/absence information observed at each sampling site via TFS and eNA analysis (Table S5); 179 species detected by TFS (Table S6); Species scores based on RDA (Table S10–S12); The abundance information at each sampling site by TFS (Table S14); Result of IndVal analysis (Table S15) (XLSX)

## AUTHOR INFORMATION

### Corresponding Authors

*Yasuaki Inoue − R&D Safety Science Research, Kao Corporation, Ichikai–Machi, Haga–Gun, Tochigi 321-3497, Japan;

*Kaede Miyata − R&D Safety Science Research, Kao Corporation, Ichikai–Machi, Haga–Gun, Tochigi 321-3497, Japan;

### Authors

Masayuki Yamane − R&D Safety Science Research, Kao Corporation, Ichikai–Machi, Haga–Gun, Tochigi 321-3497, Japan

Hiroshi Honda − R&D Safety Science Research, Kao Corporation, Ichikai–Machi, Haga–Gun, Tochigi 321-3497, Japan

### Author Contributions

Conceptualization, Y. I.; K. M. Methodology, Y. I.; K.M. Formal Analysis, Y. I. Investigation, Y. I., K.M., and H.H. Writing–Original Draft Preparation, Y.I. Writing–Review and Editing, K.M., H.H., and M.Y. Visualization, Y.I. Project Administration, H.H. and M.Y.

### Notes

The authors declare no competing interests.

## ACKNOWLEDGMENTS

We thank Mr. Takamitsu Kawaguchi, Tomohisa Nagaike, and Daiki Uchida of Bioindicator Co., Ltd., for determining the species names and classifying the habitats of the detected species. We thank Yuta Hasebe of the Kanagawa Environmental Research Center, Hiratsuka, Kanagawa, Japan, for developing the reference database.

## BRIEF

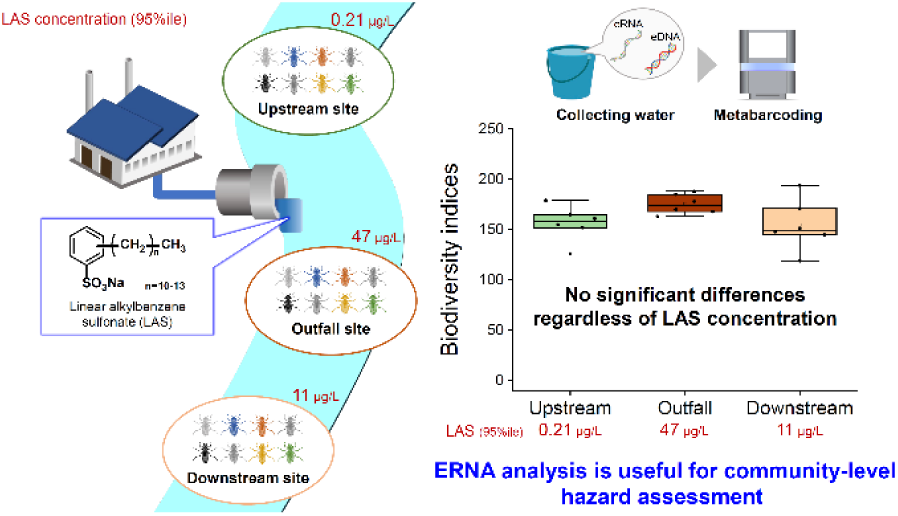

SYNOPSIS Field surveys using eRNA metabarcoding provide critical or complementary information for environmental effect assessment.

