## Supplementary Information for "Environmental RNA/DNA Metabarcoding for Ecological Risk Assessment of Chemicals Based on Response of Benthic Communities in Natural Environments"

**Contents**

1. Supplementary Texts (Text S1–3)

2. Supplementary Figures (Figure S1–2)

3. Supplementary Tables (Table S1–2, 6–8,12)

### 1. Supplementary Text

#### Text S1. Measurement of linear alkylbenzene sulfonate (LAS) concentration

LAS analysis was performed using liquid chromatography-tandem mass spectrometry (LC-MS/MS; HPLC: 1200SL, MS/MS: 6460 Triple Quad; Agilent Technologies, Santa Clara, CA, USA). C12-LAS-$^{13}C_6$ (CLM-9349-1.2; Cambridge Isotope Laboratories, Andover, MA, USA) was added to each water sample as a surrogate standard prior to column separation. Solid-phase extraction cartridges (400 mg; Sep-Pak tC18 Plus, Waters, MA, USA) were conditioned with 10 mL MeOH and 10 mL ultrapure water added at 5 mL/min. Sodium azide was added to river water and the samples passed through a cartridge at a rate of 5 mL/min. The cartridge was then cleaned with 10 mL of ultrapure water and dried for approximately 10 min under a gentle vacuum. The target compounds were eluted with 10 mL methanol and the eluate evaporated under a gentle nitrogen stream at 40 °C. Samples were reconstituted in 1 mL of a 40:60% (v/v) acetonitrile:water solution.

The mobile phase consisted of a 50 mM ammonium formate solution with 0.1% formic acid (mobile phase A) and acetonitrile (mobile phase B). Chromatographic separation was performed using a Mightysil RP-18GP column (KANTO CHEMICAL, Japan; 2.0 × 150 mm, 3 μm) at a flow rate of 0.2 mL/min using the following gradient (% mobile phase B): initial 3 min, 40%; 3 min, 40–65%; 20 min, 65%; 10 min, 95%, and 6.0 min: 40%. The column oven temperature was maintained at 40 °C and was 5 μL of sample injected. MS/MS was performed using electrospray ionization in the negative-ion mode. Analytes were acquired through multiple reaction-monitoring experiments. The monitored precursor/fragment ions were observed at m/z 297/183, 311/183, 325/183, 339/183, 353/183, and 331.2/176 for C10-LAS, C11-LAS, C12-LAS, C13-LAS, C14-LAS, and C12-LAS-$^{13}C_6$, respectively, and fragmentor voltages of 180 and 215 V were used to obtain the m/z 183 ion for C10-14 LAS and the m/z 176 ion for C12-LAS-$^{13}C_6$, respectively, with an internal standard concentration of 50 μg/L. LAS was determined using the absolute calibration curve method, with the calibration curve constructed using calibration standards at concentrations of 0.002 to 12 mg/L using the analyte/internal standard peak area ratio versus analyte concentration. The values of the coefficient of determination ($R^2$) for the calibration curves were above 0.998 for all the compounds. The limit of quantification for LAS was the lowest concentration of the standard solution (0.002 mg/L), which is within the range of the confirmed calibration. Therefore, considering the pretreatment, the limit of detection for each component and the total LAS (C10-C14 LAS) were 0.004 and 0.02 μg/L, respectively.

**Text S2. Summary of the literature used to confirm species name and habitat classification.**


1. Ogata, K. Notes on Simuliidae of the Ryukyu Islands (Diptera). *Jap. J. M. Sc. & Biol.* 1956, *9*, 59–69. DOI: 10.7883/yoken1952.9.59

2. Kawamura, T.; Ueno M. *Freshwater Biology of Japan*; Hokuryukan, Tokyo, 1973 (in Japanese).

3. Nishimura, S. *Guide to seashore animals of Japan with color pictures and keys-1*; Hoikusha, Osaka 1992 (in Japanese).

4. Nozaki, T.; Ito, T.; Tanida, K. Checklists of Trichoptera in Japan 2. Glossosomatidae, Beraeidae, Odontoceridae and Molannidae. *Jap. J. Limnol.* 1994, *55* (4), 297–305. DOI: 10.3739/rikusui.55.297

5. Nishimura, S. *Guide to seashore animals of Japan with color pictures and keys-2*; Hoikusha, Osaka, 1995 (in Japanese).

6. Kagaya, T., Nozaki, T., Kuranishi, R. and Katagiri, K., Eds. *Fauna and distribution of Trichoptera in the Tama- River system*; Tokyu Foundation for Better Environment, Tokyo, 1998 (in Japanese).

7. Nozaki, T.; Tanida, K.; Ito, T. Checklists of Trichoptera in Japan 3. Limnocentropodidae, Phryganopsychidae, Phryganeidae, Brachycentridae, and Apataniidae. *Jap. J. Limnol.* 1999, *60* (3), 347–366. DIO: 10.3739/rikusui.60.347

8. Nozaki, T.; Tanida, K.; Ito, T. Checklists of Trichoptera in Japan. 4. Goeridae, Uenoidae, and Limnephilidae. *Limnology.* 2000, *1*, 197–208. DOI: 10.1007/s102010070007

9. Kondo, S.; Hirabayashi, K.; Iwakuma, T.; R. Ueno. *The World of Chironomidae*; Baifukan, Tokyo, 2001 (in Japanese).

10. Nozaki, T. Revision of the genus Nothopsyche Banks (Trichoptera: Limnephilidae) in Japan. *Entomol. Sci*. 2002, *5 (1)*, 103–124.

11. Nishijima, S.; Nishida, M.; Shikatani, N.; Shokita, S. *The flora and fauna of inland waters in the Ryukyu islands*; Tokai University Press, Hatano, 2003 (in Japanese).

12. Masuda, O.; Uchiyama, R. *Guide to Freshwater Shellfish of Japan*; Pisces, Yokohama, 2004 (in Japanese).

13. Kondo, S.; Mano, T.; Yamamoto, M.; Kobayashi T. Chironomid midges emerged from an *Egeria densa* community at middle reaches of the Yahagi River during fall season in 2003. Yahagigawa kenkyu 2005, *9*, 49–53 (in Japanese).

14. Abe, H.; Ishii, N.; Ito, T.; Kaneko, N.; Maeda, K.; Miura, S.; Yoneda, M. *A guide to the mammals of Japan*; Tokai University Press, Hatano, 2008 (in Japanese).

15. Fujitani, T. Japanese Baetidae (Ephemeroptera): keys to seven genera with information on taxonomy, distribution and habitat. *Jap. J. Limnol.* 2006, *67*, 185–207. DOI: 10.3739/rikusui.67.185

16. Kobayashi, H.; Idei, M.; Mayama, S.; Nagumo, T.; Osada, K. *H.Kobayasi's atlas of Japanese diatoms based on electron microscopy*; Uchida Rokakuho, Tokyo, 2006 (in Japanese).

17. Kawakatsu, M.; Nishino, M.; Ohtaka, A. Currently known exotic planarians from Japan. *Jap. J. Limnol.* 2007, *68*, 461–469 (in Japanese).

18. Aoki, K. Ten species of the genus *Ameletus* (Insecta: Ephemeroptera: Ameletidae) from Nagara River and Kiso River, central Japan. *Limnology in Tokai Region of Japan* 2010, *43*, 7–15 (in Japanese).

19. Japanese Association for Chironomidae Studies. *Chironomidae of Japan*; Bun-ichi Sogo Shuppan, Tokyo, 2010 (in Japanese).

20. Toyota, K.; Seki, S.; Komai, T. *Freshwater Shrimp and crab of Japan;* Seibundo Shinkosha, Tokyo, 2014 (in Japanese).

21. Hirabayashi, K.; Yamamoto, N.; Yamamoto, Y. Massive flights of nuisance insects, *Diplocladius cultriger* (Diptera: Chironomidae), in Kamikochi Station, Shinshu University, in Myojin Area in Kamikochi, Chubu Sangaku National Park, Japan. *Pestology* 2015, *30 (2)*, 73–76. DOI: 10.24486/pestology.30.2_73

22. Aoki, J. *Pictorial keys to soil animals of Japan*; Seibundo Shinkosha, Tokai University Press, Hatano, 2015 (in Japanese).

23. Kokubun, M.; Tanaka, N.; Sakae, Y.; Kaketani, R.; Nozawa, Y.; Muratu, T.; Takizawa, H.; Kosaka, I.; Abe, K. Influence on aquatic insects caused by check dam construction. *J. Jpn. Soc. Reveget. Tech.* 2015, *41 (1)*, 271–274. DOI: 10.7211/jjsrt.41.271

24. Maruyama, H.; Hanada, S. *A field guide to Japanese aquatic insects: adults of mayflies, stoneflies and caddisflies*; Zenkoku Noson Kyokai Co,Ltd., Tokyo, 2016 (in Japanese).

25. Maruyama, H.; Takai, M. *A field guide to Japanese aquatic insects: larva of mayflies, stoneflies and caddisflies*; Zenkoku Noson Kyokai Co,Ltd., Tokyo, 2016 (in Japanese).

26. Ohtaka, A.; Kobayashi, T. Seasonal changes in the composition of aquatic macroinvertebrates in an outlet stream of Lake Oike, Tsugaru Juniko Lakes, northern Japan, with special reference to adult chironomid fauna and relationships between hydropsychid trichopterans and their parasitic chironomid, *Polypedilum kamotertium. Jap. J. Limnol.* 2016, *77*, 271–279. DOI: 10.3739/rikusui.77.271

27. Kawai, S.; Tanida, K. *Aquatic Insects of Japan: Manual with Keys and Illustrations (The second edition)*; Tokai University Press, Hatano, 2018 (in Japanese).

28. Yoshitomi, H.; Hayashi, M. *Ochthebius* (Coleoptera: Hydraenidae) Inhabiting River in Shimane Prefecture, Japan. *Bull. Hoshizaki Green Found.* 2019, *22*, 77–83.

29. Hirabayashi, K.; Saito, K.; Inoue, E., Yamamoto, Y. Long-term monitoring research of chironomid fauna (Diptera, Chironomidae) in the residential area during the winter season. *Jpn. J. Environ. Entomol. Zool.* 2019, *30 (3)*, 125–134. DOI: 10.11257/jjeez.30.125

30. Hosoya, K.; Fujita, T.; Takeuchi, H.; Kawase, S.; Uchiyama, R. *Freshwater fishes of Japan*; Yama-Kei Publishers Co., Ltd., Tokyo, 2019 (in Japanese).

31. Nakajima, J.; Hayashi, M.; Ishida, K.; Kitano, T.; Yoshitomi, H. *Aquatic coleoptera and hemiptera of Japan*; Bun-ichi Sogo Shuppan, Tokyo, 2020 (in Japanese).

32. Herpetological Society of Japan; Matsui, M.; Mori, A. *Amphibians and Reptiles of Japan*; Sunrise Publisher, Hikone, 2021 (in Japanese).

### Text S3. Discussion of reasons for the increase in the total families and taxa detected by TFS at the downstream site.

#### Statistical analyses

Cluster analyses were then conducted based on the abundance datasets of TFS using Euclidean distances to characterize the communities at each sampling site, and index values (IndVal) were measured to identify the indicator species within each group determined by cluster analysis. No species satisfied the statistical threshold ($p<0.05$) in the IndVal analysis; therefore, species with $p<0.5$ were defined as indicator species. Cluster and IndVal analyses were performed using the R package stats (function: "hclust") and labdsv (function: "indval"), respectively.

#### RESULTS AND DISCUSSION

Cluster analysis based on the abundance TFS data revealed that communities at the sampling sites could be classified into two groups: the upstream/outfall sites (Group 1), and the downstream site (Group 2) (Figure S2, Table S14). IndVal analysis indicated that the number of indicator Chironomidae species that inhabit dirty environment was greater in Group 2 (six species) than Group 1 (two species) (Table S15). Species that inhabit clean aquatic environments, such as Ephemeroptera, Plecoptera, and Trichoptera, were also selected as indicator species for Group 2. The relatively high number of *E. coli* detected downstream suggests that soil and compost from the surrounding fields may flow into the river. Therefore, as mentioned in the main text, the number of detections may increase because the downstream site is a suitable environment for species that inhabit both clean and slightly dirty environments.

### 2. Supplementary Figures

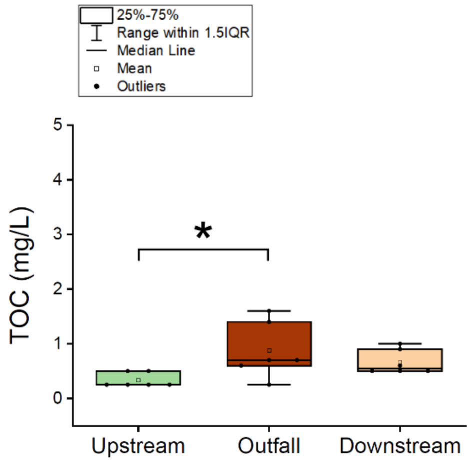


**Figure S1.** Comparison of total organic carbon (TOC) at different sampling sites. Differences between sampling sites were analyzed by Kruskal-wallis test. Asterisks indicate factors that are statistically significant following multiple comparison adjustment for Dunn's test (*: $p<0.05$).

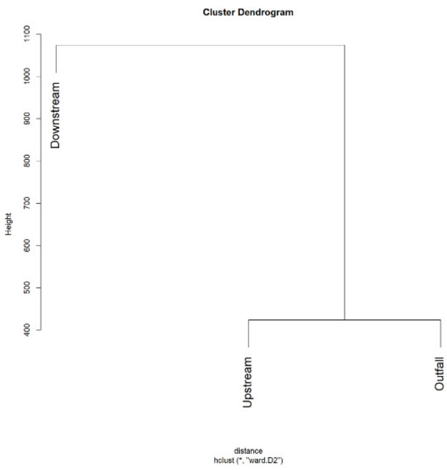


**Figure S2.** Cluster analysis results showing two groups of benthic communities. Cluster analysis was performed using the abundance data obtained via TFS and the distance matrix was calculated using a Euclidean index.

### 3. Supplementary Tables

**Table S1.** Details of field surveys and water quality measurements. Circles indicate sampling. Sampling was conducted six times at each site.

| | December 11 | December 13 | December 14 | January 15 | January 16 | January 17 | February 27 | February 28 |
|---|---|---|---|---|---|---|---|---|
| Upstream site | - | - | ○ | ○ | ○ | ○ | ○ | ○ |
| Outfall site | ○ | - | - | ○ | ○ | ○ | ○ | ○ |
| Downstream site | - | ○ | - | ○ | ○ | ○ | ○ | ○ |

**Table S2.** Universal primer pairs (MtInsects-16S primer set) used for metabarcoding.

| Primer | Oligonucleotide sequence (5’-3’) |
|---|---|
| MtInsects-16S F | ACACTCTTTCCCTACACGACGCTCTTCCGATCTNNNNNNGGACGAGAAGACCCTWTAGA |
| MtInsects-16S R | GTGACTGGAGTTCAGACGTGTGCTCTTCCGATCTNNNNNNATCCAACATCGAGGTCGCAA |

**Table S7.** Permutational analysis of variance (PERMANOVA) analysis of benthic communities at different sampling sites detected by eRNA metabarcoding analysis. Bonferroni's correction was applied to the adjusted *p*-values.

| | Df | SumsOfSqs | F.Model | $R^2$ | p. value | p. adjusted |
|---|---|---|---|---|---|---|
| Upstream vs Outfall | 1 | 0.05035 | 3.381 | 0.2526 | 0.002 | 0.006 |
| Upstream vs Downstream | 1 | 0.09231 | 5.734 | 0.3644 | 0.005 | 0.015 |
| Outfall vs Downstream | 1 | 0.05951 | 3.336 | 0.2502 | 0.033 | 0.099 |

**Table S8.** Permutational analysis of variance (PERMANOVA) analysis of benthic communities at different sampling sites detected by eDNA metabarcoding analysis. Bonferroni's correction was applied to the adjusted *p*-values.

| | Df | SumsOfSqs | F.Model | $R^2$ | p. value | p. adjusted |
|---|---|---|---|---|---|---|
| Upstream vs Outfall | 1 | 0.04746 | 2.285 | 0.1860 | 0.010 | 0.030 |
| Upstream vs Downstream | 1 | 0.08767 | 4.776 | 0.3232 | 0.004 | 0.012 |
| Outfall vs Downstream | 1 | 0.04528 | 1.785 | 0.1514 | 0.109 | 0.327 |

**Table S9.** Results of significant correlation tests for benthic communities detected by eRNA analysis and environmental factors selected using the forward selection procedure.

| Environmental factors | Df | Variance | F | p. value |
|---|---|---|---|---|
| *E. coil* | 1 | 3.324 | 1.834 | 0.014 |
| LAS | 1 | 2.870 | 1.584 | 0.032 |

**Table S13.** Results of significant correlation testing for benthic communities detected by eDNA analysis and environmental factors selected via forward selection.

| Environmental factors | Df | Variance | F | p. value |
|---|---|---|---|---|
| *E. coil* | 1 | 3.350 | 1.815 | 0.008 |
| LAS | 1 | 2.803 | 1.518 | 0.056 |
| pH | 1 | 2.732 | 1.480 | 0.044 |
